# Genome-wide association study of suicide death and polygenic prediction of clinical antecedents

**DOI:** 10.1101/234674

**Authors:** Anna R. Docherty, Andrey A. Shabalin, Emily DiBlasi, Eric Monson, Niamh Mullins, Daniel E. Adkins, Silviu-Alin Bacanu, Amanda V. Bakian, Sheila Crowell, Todd M. Darlington, Brandon Callor, Erik D. Christensen, Douglas Gray, Brooks Keeshin, Michael Klein, John S. Anderson, Leslie Jerominski, Caroline Hayward, David J. Porteous, Andrew McIntosh, Qingqin Li, Hilary Coon

## Abstract

**Objective:** Suicide death is a highly preventable, yet growing, worldwide health crisis. To date, there has been a lack of adequately powered genomic studies of suicide, with no sizeable suicide death cohorts available for study. To address this limitation, we conducted the first comprehensive genomic analysis of suicide death, using a previously unpublished suicide cohort.

**Methods:** The analysis sample consisted of 3,413 population-ascertained cases of European ancestry and 14,810 ancestrally matched controls. Analytical methods included principle components analysis for ancestral matching and adjusting for population stratification, linear mixed model genome-wide association testing (conditional on genetic relatedness matrix), gene and gene set enrichment testing, polygenic score analyses, as well as SNP heritability and genetic correlation estimation using LD score regression.

**Results:** GWAS identified two genome-wide significant loci (6 SNPs, *p*<5×10^−8^). Gene-based analyses implicated 19 genes on chromosomes 13, 15, 16, 17, and 19 (*q*<0.05). Suicide heritability was estimated *h^2^* =0.2463, SE = 0.0356 using summary statistics from a multivariate logistic GWAS adjusting for ancestry. Notably, suicide polygenic scores were robustly predictive of out of sample suicide death, as were polygenic scores for several other psychiatric disorders and psychological traits, particularly behavioral disinhibition and major depressive disorder.

**Conclusions:** In this report, we identify multiple genome-wide significant loci/genes, and demonstrate robust polygenic score prediction of suicide death case-control status, adjusting for ancestry, in independent training and test sets. Additionally, we report that suicide death cases have increased genetic risk for behavioral disinhibition, major depression, autism spectrum disorder, psychosis, and alcohol use disorder relative to controls. Results demonstrate the ability of polygenic scores to robustly, and multidimensionally, predict suicide death case-control status.

---

Suicide death is a behavioral event which reflects a complex, heritable phenotype with diverse clinical antecedents and environmental contributing factors.^1^ The rate of suicide death has been steadily increasing_2_ and, in the United States, suicide is now ranked the second leading cause of death for all persons 15-24 years old.^3^ Despite the significant heritability of suicide death,^4^ genetic research on suicide has generally been limited to the study of suicide-related behaviors, rather than the extreme phenotype of suicide death. Despite promising initial results in GWAS meta-analyses of suicidal behavior,^5^ most suicidal behavior does not result in suicide death.^1,6^ Thus, suicidal behavior likely represents a more heterogeneous and less severe phenotype than suicide death, and these features likely adversely affect statistical power to detect associated genomic signals. Conversely, the unambiguous phenotype of suicide death avoids several confounds inherent in the study of suicidal behavior or ideation, and also focuses study on one of single most critical public health outcomes in contemporary America.

Previous genetic research on suicidal behavior phenotypes has tended toward narrow ascertainment, studying only individuals with specific diagnoses (e.g., mood disorders, psychotic disorders) in order to maximize severity and accommodate post hoc study design. However, in population-based sample with ascertainment wholly independent of any co-occurring diagnoses, like the current study, the distribution, prevalence, and interaction of variables will be more representative of the corresponding population of suicide deaths, *ceteris paribus*.

Suicide death is a complex behavioral phenotype that, like its associated clinical antecedents including schizophrenia and depression, reflects a complex, and likely highly polygenic, etiology.^5,7^ Currently, the scientific literature lacks robust examination of suicide death in relation to molecular genetic risk for any medical or psychiatric diagnoses, and no polygenic scores have yet been developed for the critical outcome of suicide death. This study addresses these gaps in knowledge, leveraging the world’s largest DNA databank of suicide death, merged with a massive bank of electronic medical record and demographic data for all cases^8,9^, to comprehensively model common variant genetic, and clinical phenotypic, precursors of suicide death.

This study represents the first adequately powered genome-wide association study of suicide death. Further, analyses leveraged comprehensive data on modes of suicide death, medical and psychiatric diagnostic (ICD-10) codes,^10^ and medical and psychiatric polygenic scores to predict common variant genetic risk for suicide death. Secondary analyses of sex differences were conducted in response to the substantial sex differences in suicide death rates and modes of suicide death.^11,12^ This study achieved reliable and robust prediction of case-control status, adjusting for ancestry and sex, using both (1) a novel polygenic score for suicide death and (2) polygenic scores for a range of comorbid psychiatric and medical risk factors, particularly behavioral disinhibition and major depressive disorder. We additionally examined gene set enrichment, SNP heritability and genetic correlations in this unique, unpublished data resource. Finally, we review important considerations relating to ancestry in suicide genetics research.

## METHODS

### Sample Ascertainment

#### Cases

In collaboration with the centralized, statewide Utah Office of Medical Examiner (OME), the authors obtained DNA samples from ~6000 persons who died by suicide. The centralized OME and conservative determination helped to maximize the accuracy of suicide case status.^13^ Suicide cause-of-death determination results from a detailed investigation of the scene of the death and circumstances of death, determination of medical conditions by full autopsy, review of medical and other public records concerning the case, interviews with survivors, in addition to standard toxicology workups. Suicide determination is traditionally made quite conservatively due to its impact on surviving relatives. DNA from suicide deaths extracted from whole blood using the Qiagen Autopure LS automated DNA extractor (www.qiagen.com). Genotyping was performed on 4,381 of these cases, as described below. After quality control procedures and ancestry analysis, data comprised 3,413 Utah suicide deaths. The Utah population is primarily Northwestern European in ancestry, a relatively genetically homogeneous group with very low inbreeding across generations, comparable to the rest of the United States.^14^ Suicide determination results from a detailed investigation of the scene of the death and circumstances of death, determination of medical conditions by full autopsy, review of medical and other public records concerning the case, interviews with survivors, in addition to standard toxicology workups. Suicide determination is traditionally made quite conservatively due to its impact on surviving relatives. The centralized OME and conservative determination helped to maximize the accuracy of suicide case status.^13^

#### Controls

##### Generation Scotland

Controls closely matching the Northern European ancestry of the cases were obtained from previously curated datasets in the UK. Wave 1 analysis included 3,623 founder controls from the population-based Generation Scotland Scottish Family Health Study.^15^ The Generation Scotland Scottish Family Health Study (N > 24,000) constitutes an ancestrally comparable population-based cohort for comparison with the suicide decedents in Utah. To eliminate confounding arising from intra-dataset relatedness, only the 3,623 founders from the Generation Scotland dataset were used in analyses.

##### UK10k

A total of 11,049 UK10K controls^16^ were analyzed in wave 2 and GWAS analyses of both waves. This second control cohort is comprised of approximately 4000 genomes from the UK along with 6000 exomes from UK individuals with selected health phenotypes. We chose these data due to the extensive phenotyping and characterization of any medical conditions present, and to avoid choosing a cohort of entirely psychiatrically and medically healthy individuals. 4,000 highly phenotyped “super control” samples were supplied from the King’s College London registry and the Avon Longitudinal Study of Parents and Children. UK10K was a collaborative project to examine obesity, autism, schizophrenia, familial hypercholesterolemia, thyroid disorders, learning disabilities, ciliopathies, congenital heart disease, coloboma, neuromuscular disorders, and rare disorders including severe insulin resistance. Genotyping and sequencing procedures for UK10k have been previously described^16^ (http://www.uk10k.org) and all molecular genetic data from UK10k were filtered to the hard call variants present in our suicide death cohort prior to imputation of all cohorts simultaneously.

##### 1000 Genomes Reference Panel

The CEU population from the 1000 Genomes Project,^17^ which includes only Utah residents carefully screened for Northwestern European ancestry, was utilized as a model for excluding ancestrally discordant suicide and control samples. These CEU data were downloaded from the 1000 Genomes Project public repository. Unrelated individuals in the CEU provide a compelling, albeit small, ancestrally matched control resource (n = 99). A variety of candidate control samples were assessed via PCA for ancestral comparability to CEU and decedent data, with UK10k and Generation Scotland founder data representing the closest match.

Utah controls would be an ideal match for the suicide cases, but as with most GWAS, local controls were not readily available at the sample size needed for GWAS. CEPH ancestry 1KG were a useful comparison group to assess the likelihood that UK controls were an appropriate match for the cases. In addition (and described in more detail below) we performed a GS control-UK10K control GWAS and subsequently eliminated any SNPs from the case-control analysis that evidenced signal between the control cohorts. This was performed to minimize the possibility of false positives in the case control GWAS due to population/geographic stratification across cohorts.

### Genotyping and Quality Control

Suicide cases were genotyped using Illumina Infinium PsychArray platform measuring 593,260 single nucleotide polymorphisms (SNPs). Generation Scotland samples were genotyped using Illumina OmniExpress SNP GWAS and exome chip, measuring 700,000 and 250,000 SNPs, respectively.^15^ UK10k samples were whole genome sequenced^16^ and variants were extracted to match the available QC’d hard-called 7,519,308 variants in the suicide cases. Genotypes were subsequently imputed in all cases and controls (details of imputation are presented in Analytics, below). Both case and control datasets resulted from population-based ascertainment, and cryptic relatedness was modeled via the derivation of genomic relatedness matrices. Genotyping quality control was performed using SNP clustering in Illumina Genome Studio https://www.illumina.com/techniques/microarrays/array-data-analysis-experimentaldesign/genomestudio.html). SNPs were retained if the GenTrain score was > 0.5 and the Cluster separation score was > 0.4. SNPs were converted to HG19 plus strand, and SNPs with >5% missing genotypes were removed. Samples with a call rate < 95% were removed.

Prior to case-control GWAS, a control-control GWAS was run (using the same methods described in Analytics: GWAS, below) to detect signal between control groups to filter out of the casecontrol GWAS (control-control *q* value >.10). For example, chromosome 4 variants within the MHC are often filtered from analyses involving Scottish controls, due to prevalent population stratification in this region. We performed a stringent screen for population-specific signal in the controls. While this method is somewhat conservative, it was deemed necessary to address potential geographic stratification confounds. Signal detected from this control-control GS vs. UK10k comparison was then filtered from subsequent case-control analyses. For the purposes of future meta-GWAS analyses, and because the MHC is relevant to psychiatric risk, we included these filtered variants in a second version of our summary statistics, available upon request.

### Data analysis

#### Principal Components Analysis (PCA)

Supplementary Figure S1 shows 1000 Genomes (1KG) superpopulations and suicide case/control samples, both included and excluded, plotted by the top 2 principal components (PC). Approximately 20% of the population-based suicide cases had a significant degree of non-Northwestern European ancestry (chiefly of admixed ancestry) and were excluded from analyses. The variation explained by top 4 PCs was reduced 7.2-fold. The top 4 PCs explain 0.89% of variation before sample filtering and 0.12% of variation after filtering, if calculated on pruned genotypes. For adequate statistical power, we examined only cases of Northern European ancestry. However, it is clear from (a) and (c) that the cohort was comprised of multiple ancestries and that research on suicide death in non-European ancestries will reflect an important step beyond this first study.

PCA was performed on control, suicide, and 1000 Genomes cohorts after LD pruning at a 0.2 threshold. To exclude ancestrally heterogeneous samples, the top principal components (defined as those components which accounted for > 0.1% of the genotype variance, *n*pc = 4) were used to establish PC centroid limits centered around 1000 Genomes CEU data, such that 99% of the CEU data fell within the limits. Only suicide and control samples also falling within these limits were considered ancestrally homogenous and thus were included in the association study. The ancestry PCA was performed using RaMWAS,^18,19^ a Bioconductor package, written by our analytical team, which comprises a complete toolset for high dimensional genomic analyses. RaMWAS includes functions for PCA for capturing batch effects and detection of outliers, association analysis while correcting for top PCs and covariates, creation of QQ-plots and Manhattan plots, and annotation of significant results.

#### Imputation

European ancestry cases and controls were well-matched to 1000 Genomes CEPH. The Haplotype Reference Consortium is comprised in part by UK controls used here, so we elected to impute genotypes based on the 1000 Genomes reference panel using minimac3^20^ and Eagle.^21^ SNPs with ambiguous strand orientation, >5% missing calls, or Hardy-Weinberg equilibrium p < 0.001 were excluded. SNPs with minor allele frequency below 0.01 or imputation R^2^ < 0.5 were also excluded. Genomic data were handled using PLINK. ^22,23^ Final GWAS analysis was performed on 7,519,308 variants passing quality control.

#### Genome-wide Association Testing

A Linear Mixed Model (LMM) algorithm tested variant association with suicide death, with follow-up examination of significant hits for linkage disequilibrium and gene set enrichment. GWAS were performed using GEMMA,^24^ a computationally efficient and open-source LMM algorithm for GWAS that models population stratification remaining after PCA by use of genomic relatedness matrices. Sex was not included as a covariate in GWAS analyses due to the association of suicide with sex status at a ratio of approximately 3:1 males: females. GWAS with hard call-only and then with imputed data were examined separately to assess potential population stratification unique to our imputed GWAS. Prior to case-control GWAS, control-control GWAS was implemented to filter signal likely due to population stratification in the controls.

##### Power Analysis

With a suicide death prevalence rate of approximately.002%, this GWAS of European ancestry with 3,413 population-based suicide deaths and 14,810 ancestry-matched controls would be expected to have at least 80% power to detect common variants (MAF ≥ 0.15) with effect sizes ≥ 1.20 at *P*<5×10^−8^ and *P*<1×10^−6^ (Supplementary Figure S2). Power at *P* <□1□×□10^−6^ is relevant because 52 SNPs reach that threshold in the current analysis. Power is lower for less-common variants and in secondary analyses stratifying by mode of suicide death and sex.

#### Gene and Gene Set Enrichment and Functional Mapping

SNP results from the GWAS were then mapped to genes within 1kb of the SNP and these genes were examined for gene set enrichment and LD using FUMA^25^ and GREAT.^26^ FUMA annotates SNPs, uses MAGMA to identify associated genes (of approximately 18,612) and provide gene and gene pathway enrichment analysis (of approximately 10,649 pathways). GREAT analyzes the functional significance of sets of *cis*-regulatory regions, by modeling the genome regulatory landscape using multiple information sources and can determine the functional domain of intergenic variants. GREAT improves upon the identification of genes associated with non-coding genomic regions through statistically rigorous incorporation of more distal binding sites from 20 ontologies. The GWAS catalog (https://www.ebi.ac.uk/gwas/) includes studies and associations if they: include a primary GWAS analysis from >100,000 SNPs, SNP-trait p-value <1.0 × 10-5 in the overall (initial GWAS + replication) population. The most significant SNP from each independent locus is extracted.

#### Polygenic Risk Scores, SNP Heritability (*h*^2^), and Genetic Correlations (*r*_G_)

Discovery GWAS summary statistics for phenotypes were compiled to score each cohort for polygenic risk. PS for suicide death was derived using PRSice 2.0^27^ and summary statistics from a 10-fold cross validation procedure to avoid overfitting. To elaborate, k-folds cross-validation is a well-established method to allow out-of-sample prediction,^28^ allowing a single dataset to unbiased serve as both training and testing data, for the purpose of suicide death polygenic score development and validation. We conservatively set the p-value threshold for predicting case status based on the data to 1.0. This eliminated overfitting arising from choosing the threshold based on the phenotype.

Using related methods, we calculated polygenic scores for several psychiatric and psychological traits in the current dataset. Of several thousand medical and psychological GWAS now available, only GWAS with N>10,000 individuals and >1,000 cases (or for population-based studies, adequate base rates) were selected for these analyses. These generally included the largest medical and psychiatric GWAS and when several versions of GWAS were available for the same phenotype (for example, neuroticism or depression) we selected the most comprehensive. For a helpful reference to GWAS available, see Watanabe et al.’s online GWAS Atlas (http://atlas.ctglab.nl/). PRSice 2.0 was used to calculate individual PS for 59 phenotypes with estimated risk allele effect sizes for each discovery sample trait. A PS is traditionally calculated as a weighted sum score, where a score for an individual in the target sample is calculated by the summation of each SNP multiplied by the effect size of that SNP in the discovery GWAS. Based on the cross-disorder psychiatric genomics findings to date, we hypothesized significant positive prediction of suicide with PS for depressive symptoms, depressive disorders, behavioral disinhibition, schizophrenia, autism, loneliness, child IQ, alcohol use, and neuroticism. Betas, p-values, and q-values (after correcting the p-values for the FDR of 5%) for association of all polygenic scores, adjusting for five ancestry principle components, to suicide death are presented in Supplementary Tables.

LDSC was used to calculate common variant *h*^2^ using summary statistics from a logistic regression model with five ancestry covariates and pruning related samples at 0.2 *p*□ from IBD. LDSC was also used to calculate common variant molecular genetic correlations (*r*_G_) with psychiatric and medical phenotypes. Finally, we performed secondary analyses to characterize genetic predictors, and clinical antecedents of (Supplementary Methods S1), mode of suicide death (Supplementary Methods S2). Specifically, in these secondary analyses, we conducted (1) epidemiological association tests between four sufficiently prevalent/powered modes of death (i.e., gun, overdose, asphyxiation, and violent trauma) and 30 ICD-10 derived clinical antecedents, and (2) association tests the suicide polygenic score with modes of death, adjusting for five ancestry covariates in multivariate regressions.

#### Sex Differences

As suicide rates and modes of suicide death are characterized by substantial sex differences^11,12^, we performed secondary epidemiological and genomic analyses to characterize sex differences in mode of death and clinical antecedents. These sex stratified analyses mirrored the full sample analyses described above, including (1) sex stratified epidemiological association tests between four sufficiently prevalent/powered modes of death (i.e., gun, overdose, asphyxiation, and violent trauma) and 30 ICD-10 derived clinical antecedents, and (2) sex stratified association tests the suicide polygenic score with modes of death, adjusting for five ancestry covariates in multivariate regressions. We constrained these exploratory analyses to only those medical diagnoses with frequencies high enough in either females or males to provide decent power for testing and report false discovery rate (FDR) corrected p-values; nonetheless, it is worth noting that power for these secondary analyses of sex differences was limited by N, which was restricted to cases-only and stratified by sex and mode of death.

## RESULTS

### Genome-wide Association

A total of six variants from two loci met genome-wide criteria for statistical association with suicide death (p < 5×10^−8^). An additional 52 variants were “nominally significant” at q < 0.1 and mapped to 19 genes. (λ = 1.015, Figure 1 and Table 1).^29,30^ All results on the full cohort are derived from analyses adjusting for effects of ancestry and sex. Genes associated with top genomic regions are presented in Supplementary Table S1. Chromosome 13 and 15 regions were supported by additional positive results that were suggestive but below threshold. Ten additional genes were identified in gene-based tests meeting threshold for nominal significance (Figure 2). The large number of signals in the SNP-based tests prompted quality control analyses varying the degree of LD pruning prior to PCA for the purposes of sensitivity analysis, and results and respective λ’s were consistent across these analyses. Supplementary Figures S3-S10 present additional plots of the top signals in each of nine regions.

**Figure 1.**
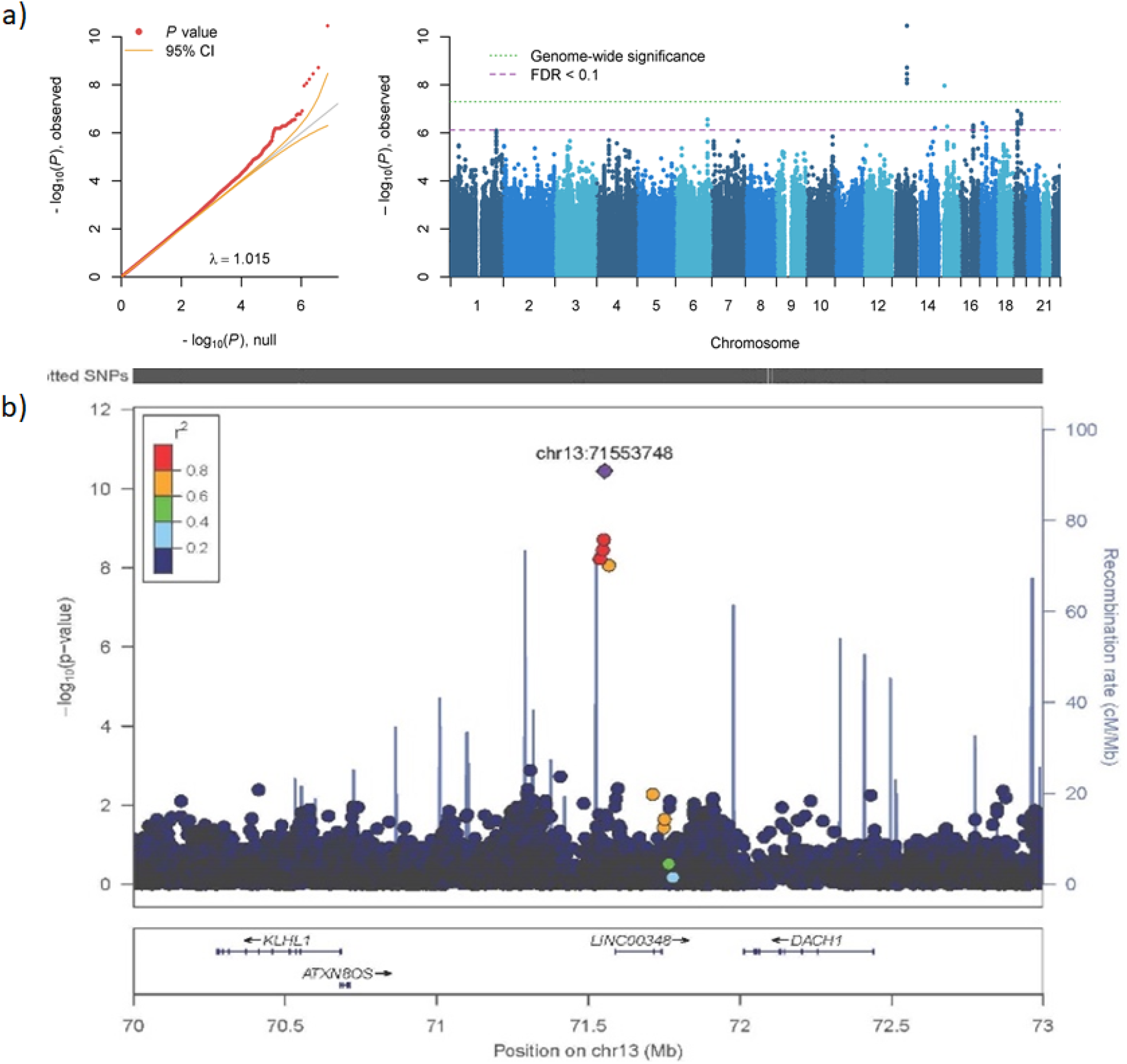
GWAS Results. **(a)** QQ and Manhattan plots from GWAS of suicide death. Y-axes for both plots reflect observed p-values. The x-axis on the qq-plot is the number of significant p-values expected under *H*_0_, and the x-axis on the Manhattan plot maps each chromosome. The purple dashed line indicates threshold for false discovery rate (FDR) corrected nominal statistical significance, the green dotted line representing the threshold for genome-wide significance after multiple testing (Bonferroni) correction. 57 SNPs met threshold for nominal significance and 6 met genome-wide significance. **(b)** Regional plot of genome-wide significant loci on chromosome 13.

**Figure 2.**
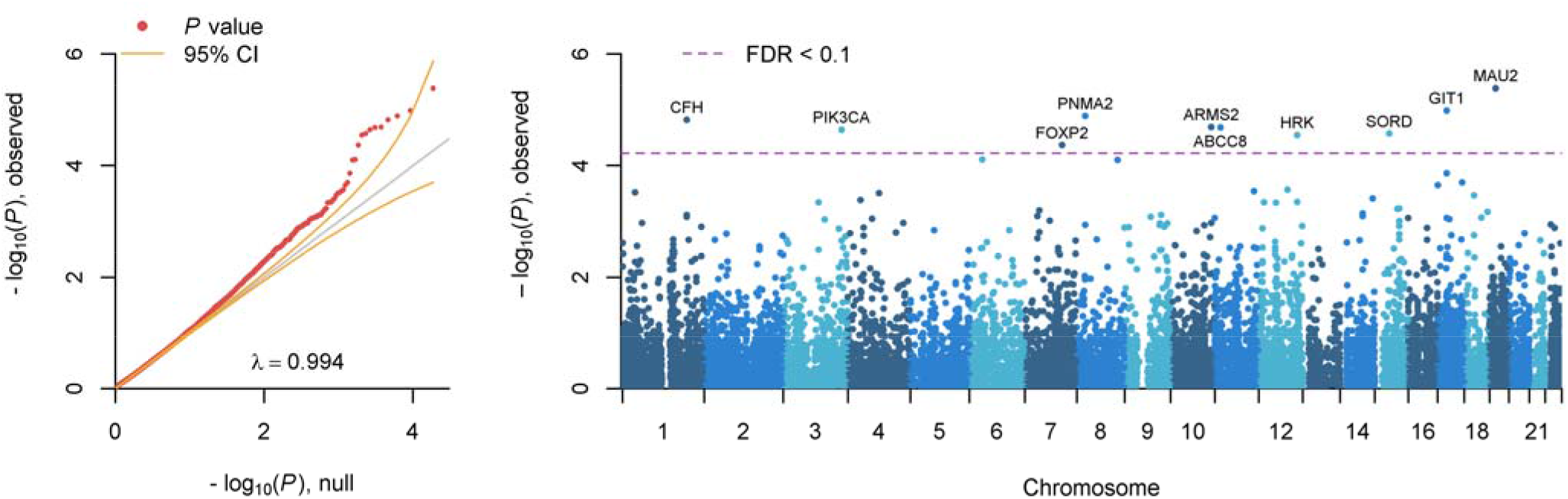
GWAS Gene-Based Results. FUMA gene-based test results: qq plot and Manhattan plot of >18,000 genes. Y-axes for both plots are identical and reflect observed p-values. The x-axis on the qq-plot is the number of significant p-values expected under *H*_0_, and the x-axis on the Manhattan plot maps each chromosome. The purple dashed line indicates threshold for FDR-corrected nominal statistical significance; 10 genes met this threshold for nominal significance.

**Table 1.** Genome-wide significant loci from GWAS of death by suicide.

| Chr | Position | SNP | A1 | A2 | AF | $\beta$ | SE | *p* value | *q* value | Nearest Gene | Function | CADD |
| --- | --- | --- | --- | --- | --- | --- | --- | --- | --- | --- | --- | --- |
| 13 | 71,538,274 | rs35502061 | G | A | 0.016 | 0.089 | 0.0153 | $5.97 \times 10^{-9}$ | $5.63 \times 10^{-3}$ | *SOGA2P1* | Intergenic | 2.69 |
| 13 | 71,547,393 | rs34053895 | A | C | 0.016 | 0.091 | 0.0154 | $3.51 \times 10^{-9}$ | $3.78 \times 10^{-3}$ | *LINC00348* | Intergenic | 0.39 |
| 13 | 71,550,518 | rs35518298 | T | C | 0.016 | 0.092 | 0.0154 | $1.92 \times 10^{-9}$ | $2.42 \times 10^{-3}$ | *LINC00348* | Intergenic | 0.33 |
| 13 | 71,553,748 | rs34399104 | T | C | 0.017 | 0.098 | 0.0147 | $3.54 \times 10^{-11}$ | $6.67 \times 10^{-5}$ | *LINC00348* | Intergenic | 19.86 |
| 13 | 71,567,365 | rs66828456 | A | C | 0.017 | 0.086 | 0.0150 | $8.63 \times 10^{-9}$ | $7.24 \times 10^{-3}$ | *LINC00348* | Intergenic | 4.24 |
| 15 | 25,962,209 | rs35256367 | G | A | 0.016 | 0.088 | 0.0155 | $1.10 \times 10^{-8}$ | $8.33 \times 10^{-3}$ | *ATP10A* | Intronic | 4.41 |

Note: SNP = single nucleotide polymorphism, CHR = chromosome, A1, A2 = alleles 1 (minor) and 2, AF = allele 1 frequency,
CADD<sup>40,41</sup> = combined annotation dependent depletion score.

### Gene and Pathway Functional Enrichment Tests

Gene-base analysis using MAGMA (FUMA^1^) identified 10 genes significantly associated (q<0.1) with suicide death (Supplementary Table S2). Additionally, mapping top SNP hits to genes suggested another 19 gene association, including chromosome 13 genes, Daschund family transcription factor 1 (*DACH1*), Ubiquitin-protein ligase protein (*UBE3A*), and Kelch-like family member 1 (*KLHL1*) on chromosome 15. Eleven of the 19 associated genes carry prior evidence of association with suicidal behaviors (Supplementary Table S2). GO pathway results included enrichment of histone modification sites *SETD6, COPR5, GATAD2A*. Full gene and Gene Ontology (GO, http://www.geneontology.org/) pathway enrichment results are presented in Supplementary Tables S3-S4. In addition to functional pathways, significant enrichment was indicated for schizophrenia results in the GWAS Catalog (*p*=1×10^−11^) (https://www.ebi.ac.uk/gwas/). Psychiatric associated traits are in green (Supplementary Table S3). IW-scoring in SNP-Nexus^31^ suggested regulatory functional significance for one SNP (chr13:71553748:C/T). Ten of the implicated genes from positional or gene-based testing have evidenced genome-wide significant differential gene expression in postmortem brain in either schizophrenia, autism, or bipolar disorder (FDR<0.1; PsychENCODE Consortium, Supplementary Table S5).^32^

### Polygenic Scores, SNP Heritability, and Genetic Correlations

In European ancestry training and test samples comprising independent case and control cohorts, and accounting for five ancestry PCs and sex, suicide PS robustly predicted suicide death case status. Suicide waves 1 and 2 comprise approximately 1,321 and 2,092 suicide cases, respectively. These predictions are plotted across 1000 p-value thresholds in Figure 3.

**Figure 3.**
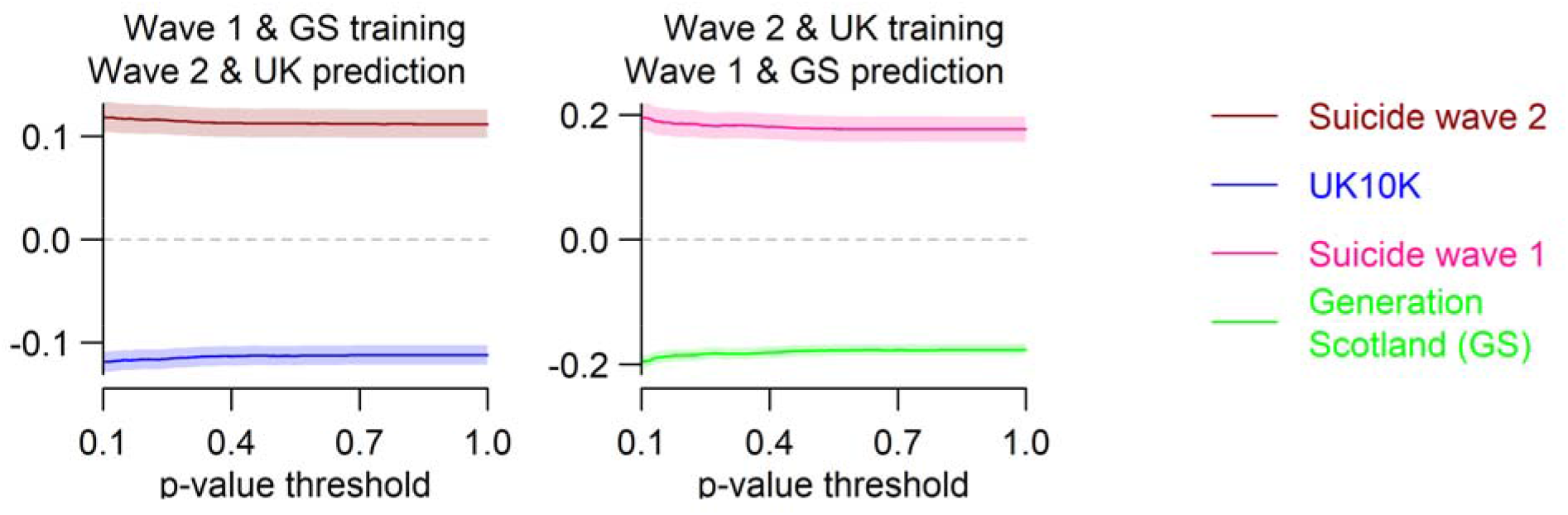
Cross Validation of Suicide Polygenic Case-Control Prediction. Polygenic prediction of suicide death case status across two independent cohorts of cases and controls. Training GWAS summary statistics are used to score the test set for suicide polygenic risk. P-value thresholds are plotted on the x-axis from 0.1-1.0, reflecting the top 10% to 100% of the common variants from the training GWAS. Standardized suicide PS scores are plotted on the y-axis. 95% confidence intervals around the scores are pictured for each cohort across all p-value thresholds.

A Linkage Disequilibrium SCore regression (LDSC)^33,34^ common variant *h^2^* estimate based on only the summary statistics from a logistic GWAS, with five ancestry covariates and pruning to remove related samples, was 0.2463, SE = 0.0356. Lambda in the latter model was inflated at 1.239. The suicide death cases differed significantly from two UK control groups on PS of phenotypes relevant to suicide death. These differences were in the expected directions. Original discovery GWAS for all phenotypes were filtered to exclude any using these control cohorts (Supplementary Table S6).

Consistent with hypotheses, significant PS elevations included alcohol use, autism spectrum disorder, child IQ, depressive symptoms, disinhibition, loneliness, and neuroticism (Figure 4). Effect sizes are communicated from the y-axis, indicating the largest effects for behavioral disinhibition and major depressive disorder PS. LD Hub^33^ provided estimates of SNP-based shared genetic covariance for several phenotypes (Supplementary Table S7). As sensitivity analysis, we disaggregated suicide by mode of death into four categories (see Supplementary Methods S2), gun, overdose, asphyxiation, violent trauma, and epidemiologically characterized these groups by association testing with 30 ICD-10 derived clinical antecedents (Supplementary Figure S11; Supplementary Tables S8-S10). Additionally, we conducted PS association testing to mode of death in all cases and no associations met multiple testing adjusted significance criteria (q<0.1; Supplementary Figure S12; Supplementary Tables S11-S13).

**Figure 4.**
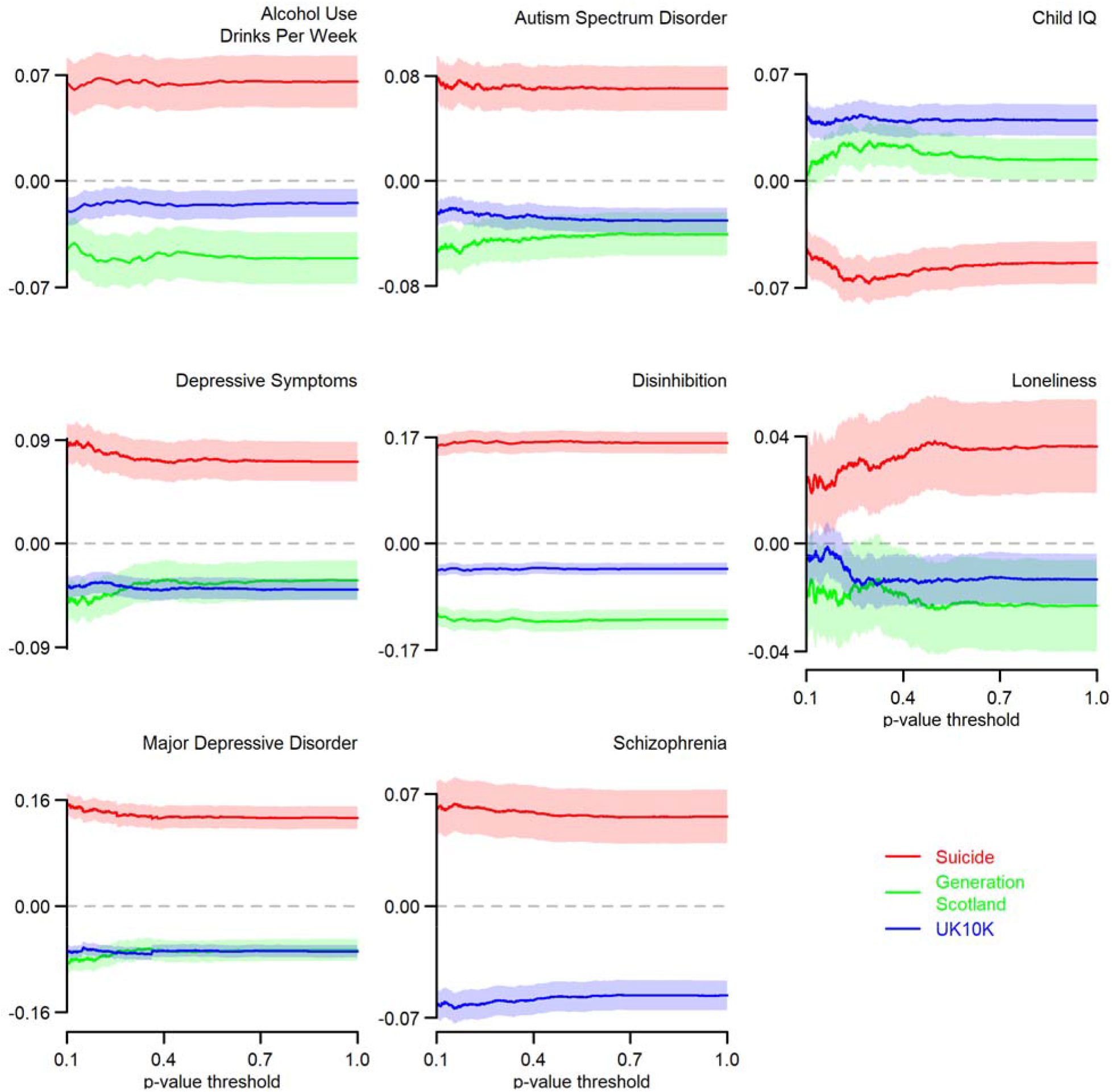
Notable Elevations of Psychiatric Polygenic Risk in Suicide Cases. Standardized polygenic scores for eight phenotypes hypothesized to be relevant to suicide (y-axis) plotted for suicide death case, and GS and UK10k control groups across a broad spectrum of PS p-value thresholding. P-value thresholds are plotted on the x-axis from 0.1-1.0. 95% confidence intervals around the scores are pictured for each cohort across p-value thresholds. Largest effect sizes, and significance levels, were observed for behavioral disinhibition and major depressive disorder.

### Sex Differences

Epidemiological analyses of sex differences indicated that suicide cases of both sexes evidenced clinical diagnostic clusters of 1) internalizing-trauma-cluster B psychiatric disorders and 2) metabolic-cardiovascular-obesity medical disorders (Supplementary Tables S14-S19). Female cases were observed to have a higher overall number of diagnoses relative to males, which could reflect increased severity in females, decreased severity in males, decreased likelihood of males receiving diagnosis, and/or decreased help-seeking in males. This is broadly consistent with observed higher relative prevalence rates of gun-related death in males and overdose death in females. All associations of PS with mode of death are presented for females and males separately in Supplementary Figures S15-S16. All PS analyses include ancestry covariates. All corresponding statistics are reported in Supplementary Tables S20-S25. No sex stratified polygenic scores findings met multiple testing adjusted significance criteria (q<0.1).

## DISCUSSION

Results from this analysis, the first adequately powered comprehensive genomic study of suicide death, yield several insights into the process and suggest important applications for future extension. The GWAS of suicide death identified two genome-wide significant loci on chromosomes 13 and 15. Moreover, 11 of the 19 genes implicated by top GWAS hits overlap with schizophrenia results from the GWAS Catalogue, and two of these 11 genes have prior associations with risk of suicidal behavior (*HS3ST3B, NCAN;* for relevant literature see Supplementary Table S26). Additionally, using k-folds cross-validation methods we were able to robustly predict suicide death case-control status using polygenic scores in out-of-sample prediction. Perhaps most compellingly, we found suicide death was strongly associated with polygenic scores for multiple psychiatric and psychological traits. These included alcohol use, autism spectrum disorder, child IQ, depressive symptoms, behavioral disinhibition, and loneliness. These results were notably consistent with expectations, as the strongest associations were to behavioral disinhibition and major depressive disorder, arguably the two most critical antecedents of suicide death^35^.

Genetic overlap of actual suicide death with suicidal behaviors remains unclear to date. More common suicidal behaviors are difficult to quantify, do not effectively predict suicide mortality, and represent individuals with a range of risk for later suicide.^36^ Moving closer to developing objective risk measures of suicide risk, future modeling of the shared genetic covariance of suicide death and suicide behaviors may isolate important genetic and environmental moderators of risk of death. Further, as genomic prediction will likely improve in the near term, as a function of increased sample sizes in suicide death cohorts, we will soon face both technical opportunities and ethical challenges associated with integrating genomic risk prediction with machine learning predictive models of suicide risk.^37,38^

Importantly, moving forward we must prioritize study genetic risk in other ancestry groups to address the potential for increasing health disparities stemming from polygenic risk research that relies only on European ancestry summary statistics.^39^ To this end, the authors are working toward cross-ancestry replication of results in individuals of Mexican American ancestry with ongoing collection of population-based cases. Future priorities also include analyses of structural variation, methylation, and predicted gene expression in suicide, investigation of the potential mediating role of substance/alcohol use in suicide, and use of genetic risk metrics to enhance predictive models of suicide.

## Acknowledgements

This work was supported by the National Institute of Mental Health (H.C., grant number R01MH099134; A.D., grant number K01MH093731), a research contract from Janssen Research, LLC (H.C. & Q.L.); the American Foundation for Suicide Prevention (A.D. & A.B.), the Brain & Behavior Research Foundation (A.D., NARSAD Young Investigator Award); the University of Utah EDGE Scholar Program (A.D.); the Clark Tanner Foundation (HC, TD, AVB), and a donation from the Sharon Kae Lehr Endowed Research Fund in Memory of James Raymond Crump (DG). Partial support for all datasets housed within the Utah Population Data Base is provided by the Huntsman Cancer Institute (HCI), http://www.huntsmancancer.org/, and the HCI Cancer Center Support grant, P30-CA-42014 from the NIH. Generation Scotland received core support from the Chief Scientist Office of the Scottish Government Health Directorates (CZD/16/6) and the Scottish Funding Council (HR03006). Genotyping of the GS samples was carried out by the Genetics Core Laboratory at the Wellcome Trust Clinical Research Facility, Edinburgh, Scotland and was funded by the Medical Research Council UK and the Wellcome Trust (Wellcome Trust Strategic Award “Stratifying Resilience and Depression Longitudinally” (STRADL) Reference 104036/Z/14/Z). Genotyping was performed by U of Utah Genomic Core and by Illumina, Inc with support from Janssen Research & Development, LLC. This work was also supported by research funding from Janssen Research & Development, LLC to University of Utah.

## SUPPLEMENTARY MATERIAL

### Supplementary Methods

S1. International Classification of Diseases (ICD-10) Code Ascertainment

S2. Clinical Diagnostic Associations with Mode of Death

### Supplementary Tables

S1. Genes Associated with Top Genomic Regions

S2-S4. FUMA and GREAT Results, GWAS Catalogue, SNP Nexus

S5. Genome-wide significant differential gene expression in PsychENCODE postmortem brain in schizophrenia, autism, or bipolar disorder

S6. Polygenic Score (PS) Discovery GWAS Summary Statistic Citation Table

S7. LD Hub Genetic Correlation Results

S8-10. All Case Diagnostic Antecedents (ICD) by Mode of Suicide (MD) Association Summary Statistics: Betas, p-values, result summaries

S11-13. All Case PS-MD Summary Statistics: Betas, p-values, result summaries

S14-19. Separate Female and Male ICD-MD Summary Statistics: Betas, p-values, result summaries

S20-S25. Separate Female and Male PS-MD Summary Statistics: Betas, p-values, result summaries

S26. Gene Literature (brief)

### Supplementary Figures

S1. PCA of 1KG super-populations, cases, controls, and excluded samples

S2. Effect sizes across frequencies required to detect both genome-wide and “nominal” levels of significance

S3. Regional Plot of Chromosome 6 Region

S4. Regional Plot of Chromosome 14 Region

S5. Regional Plot of Chromosome 15 Region

S6. Regional Plot of Chromosome 15 Region

S7. Regional Plot of Chromosome 16 Region

S8. Regional Plot of Chromosome 17 Region

S9. Regional Plot of Chromosome 17 Region

S10. Regional Plot of Chromosome 19 Region

S11. Diagnostic Antecedents (ICD) of Specific Mode of Suicide (MD) Plot: All Suicide Deaths

S12. PS × MD Plot: All Suicide Deaths

S13. ICD × MD Plot: Female Suicide Deaths > 25 Years of Age

S14. ICD × MD Plot: Male Suicide Deaths > 25 Years of Age

S15. PS × MD Plot: Female Suicide Deaths Plot

S16. PS × MD Plot: Male Suicide Deaths Plot

## Notes

#### Summary of Updates

Figures

