## Supplemental Methods for "Genome-wide association study of suicide death and polygenic prediction of clinical antecedents"

**SUPPLEMENTARY METHODS**

**S1. International Classification of Diseases (ICD-10) Code Ascertainment.** Decedent data are linked within the Utah Population Database (UPDB), a unique resource that houses data on > 9 million individuals and contains vital statistic and demographic records and electronic health records from the two major hospital systems in the Utah. ICD codes for suicide cases were recovered from the electronic data warehouses of Utah’s two largest health care providers, Intermountain Healthcare and the University of Utah. Ambulatory and inpatient ICD codes were obtained directly from UPDB. EMR data, while comprehensive and population-based, are also subject to missingness (random and non-random), and severity of health disparity may correspond with the presence and absence of ICD codes in an EMR. For this reason, we checked ICD code associations with polygenic risk for the same disorders and assessed relationships of ICD with PRS across age group and date of code. The rationale for the latter analyses was that younger individuals are less likely to have specific ICD codes due to either lack of contact with the system or to lack of maturation of overt psychopathology (e.g., psychosis, personality disorders). In addition, ICD codes from early EMR development in Utah (pre-year 2000) are notably sparser. To manage the wide diversity of codes, we limited our analyses to 30 categories of codes reflecting 30 relevant diagnostic domains. Data were subsequently completely de-identified prior to analysis. The study was approved by the Institutional Review Boards of the University of Utah, Intermountain Healthcare, and the Utah Department of Health.

**S2. Clinical Diagnostic Associations with Mode of Death** are presented in Supplementary Figure S11. Suicide by gun was associated with the general absence of clinical diagnoses. Suicide by overdose was associated with many clinical diagnoses and most robustly with obesity and sleep disorders. Suicide by violent trauma was associated with a clinical diagnosis of schizophrenia. Corresponding βs and p-values for Supplementary Figure S11 are presented in Supplementary Tables S8-S10.
