## Supplemental Figures for "Genome-wide association study of suicide death and polygenic prediction of clinical antecedents"

**SUPPLEMENTARY FIGURES**

S1. PCA of 1KG super-populations, cases, controls, and excluded samples

S2. Effect sizes across frequencies required to detect both genome-wide and “nominal” levels of significance

S3. Regional Plot of Chromosome 6 Region

S4. Regional Plot of Chromosome 14 Region

S5. Regional Plot of Chromosome 15 Region

S6. Regional Plot of Chromosome 15 Region

S7. Regional Plot of Chromosome 16 Region

S8. Regional Plot of Chromosome 17 Region

S9. Regional Plot of Chromosome 17 Region

S10. Regional Plot of Chromosome 19 Region

S11. Diagnostic Antecedents (ICD) of Specific Mode of Suicide (MD)

S12. PRS x MD: All Suicide Deaths

S13. ICD x MD plots: Female suicide deaths > 25 years of age

S14. ICD x MD plots: Male suicide deaths > 25 years of age

S15. PRS X MD: Female Suicide Deaths

S16. PRS X MD: Male Suicide Deaths

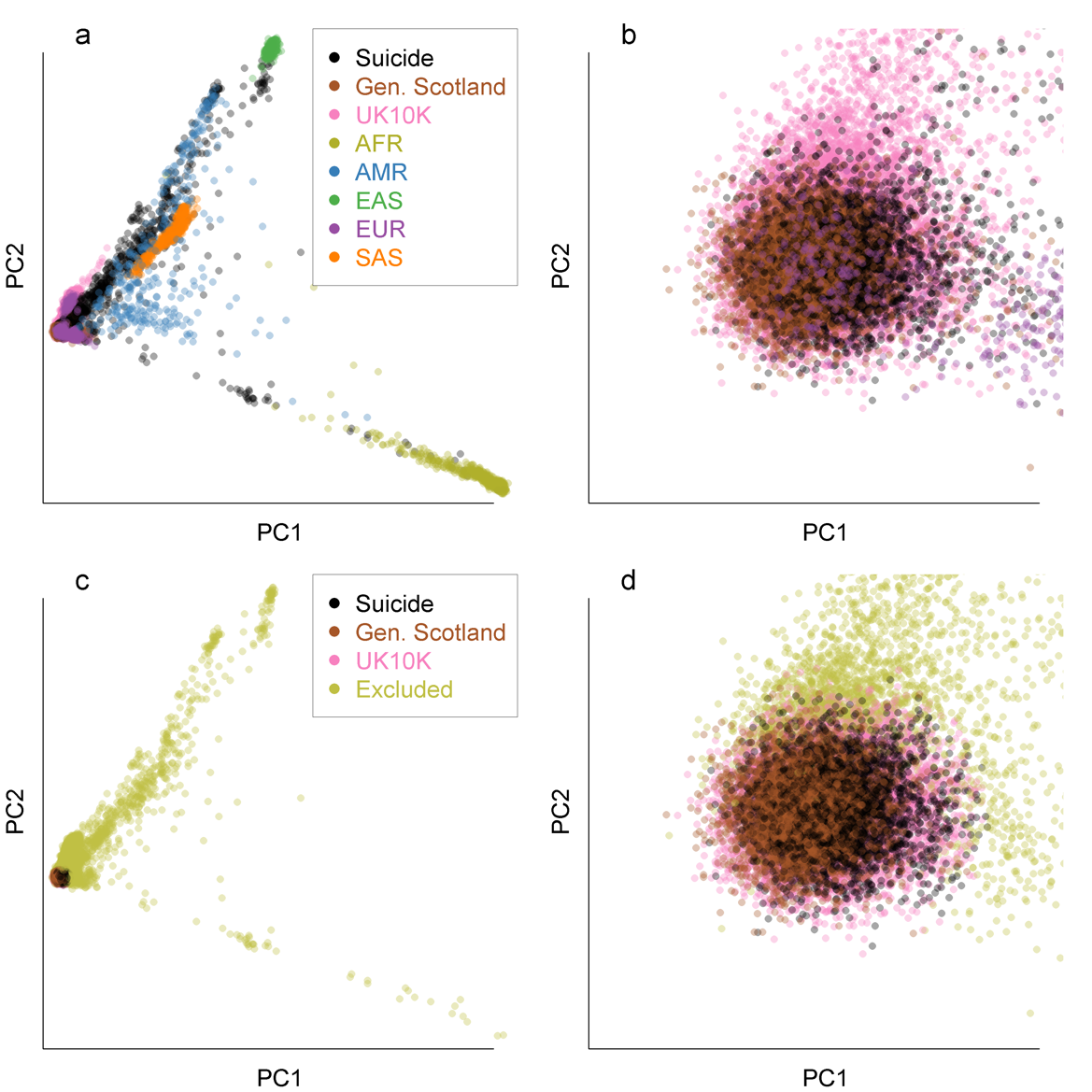

**S1. PCA of 1KG super-populations, cases, controls, and excluded samples**. Suicide death samples plotted by the principal component explaining the most variance (PC1) versus the principal component explaining the second most variance (PC2) in all 1KG super-populations (a) and focusing on only the Northern European cases and controls (b). (c) and (d) highlight excluded cases. For adequate statistical power, we examined only cases of Northern European ancestry. However, it is clear from (a) and (c) that the cohort was comprised of multiple ancestries and that research on suicide death in non-European ancestries will reflect an important step beyond this first study.

**S2. Power Plot: Power to detect both genome-wide and nominal levels of significance across MAF.**

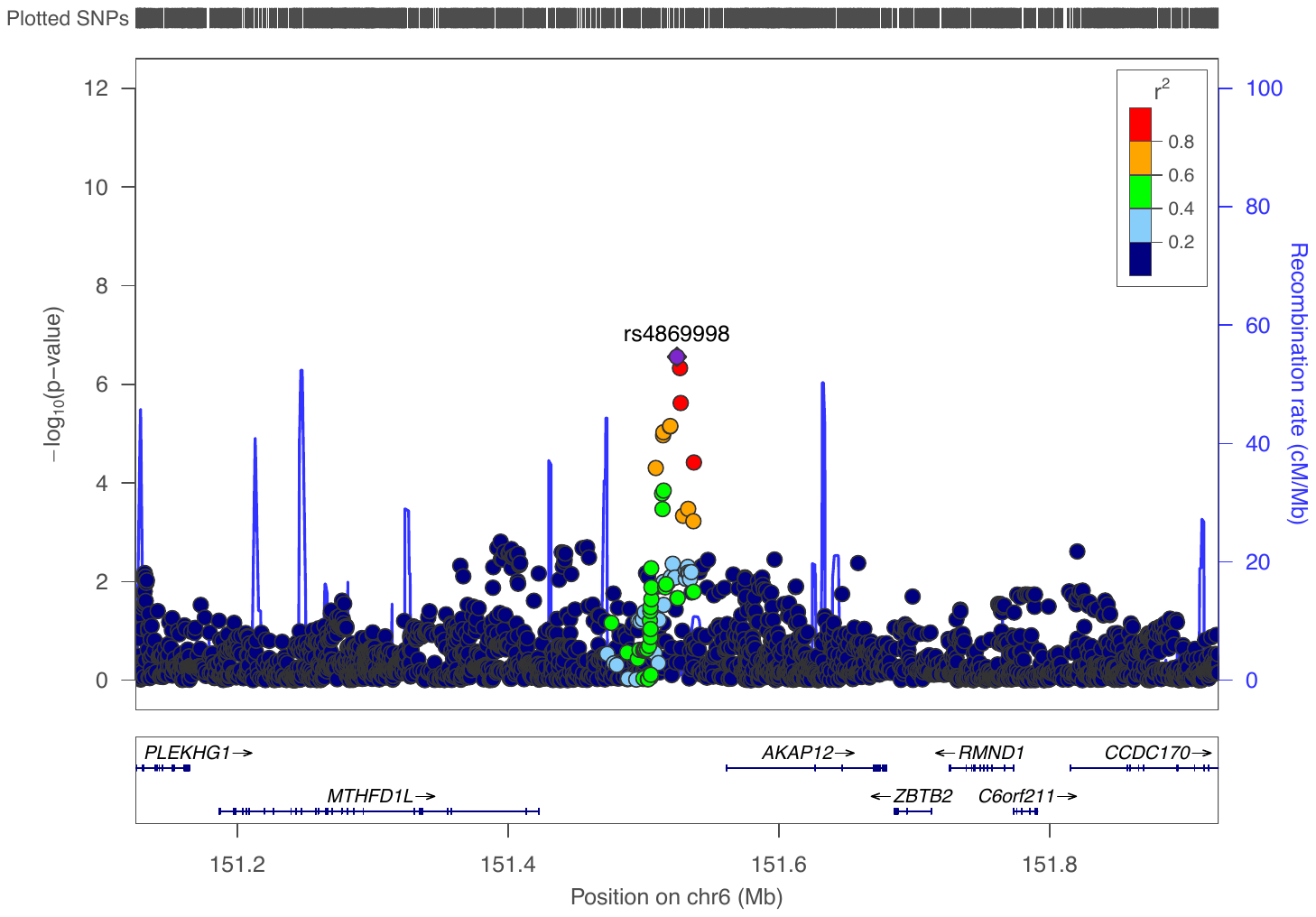

**S3. Chromosome 6**

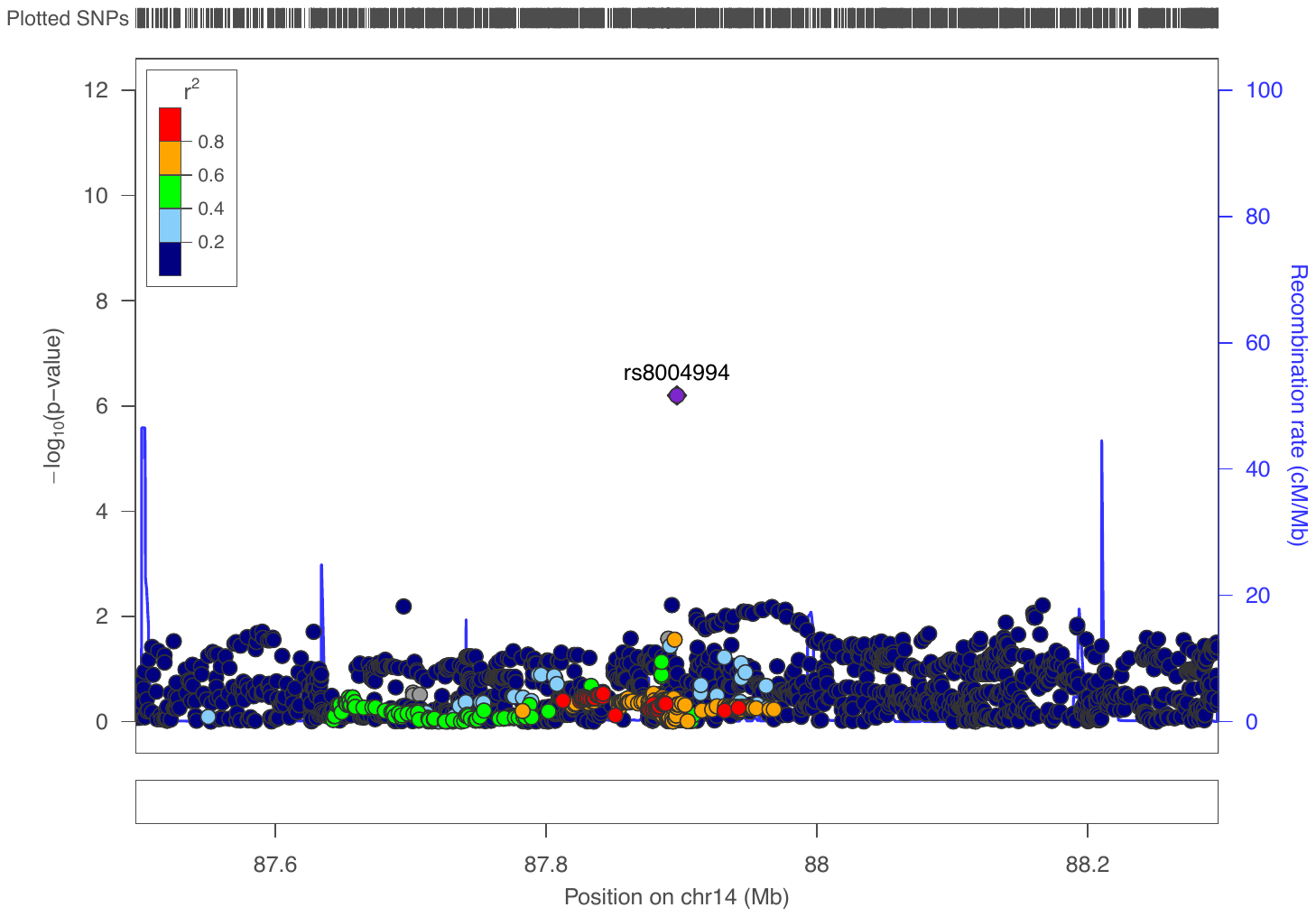

**S4. Chromosome 14**

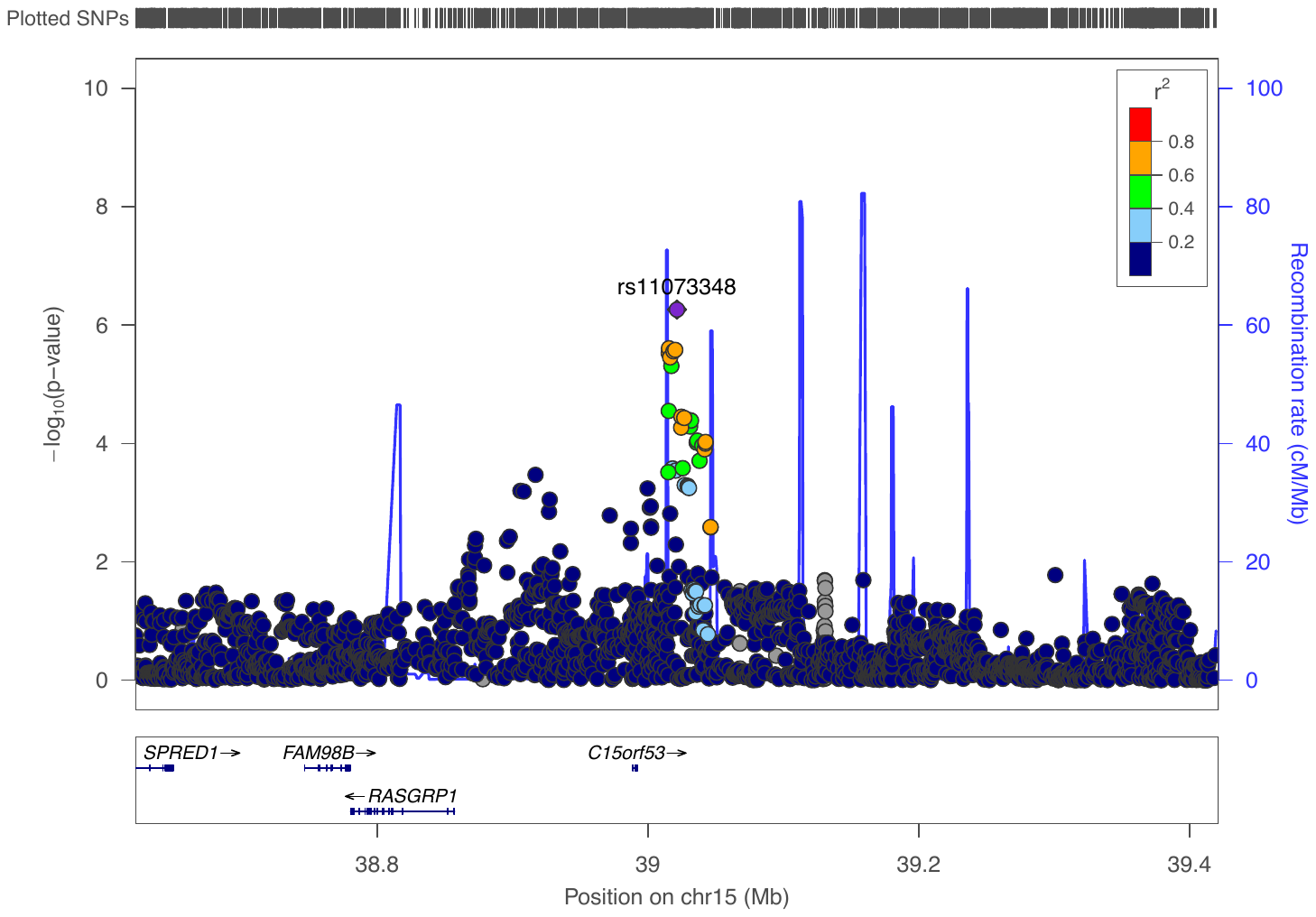

**S5. Chromosome 15**

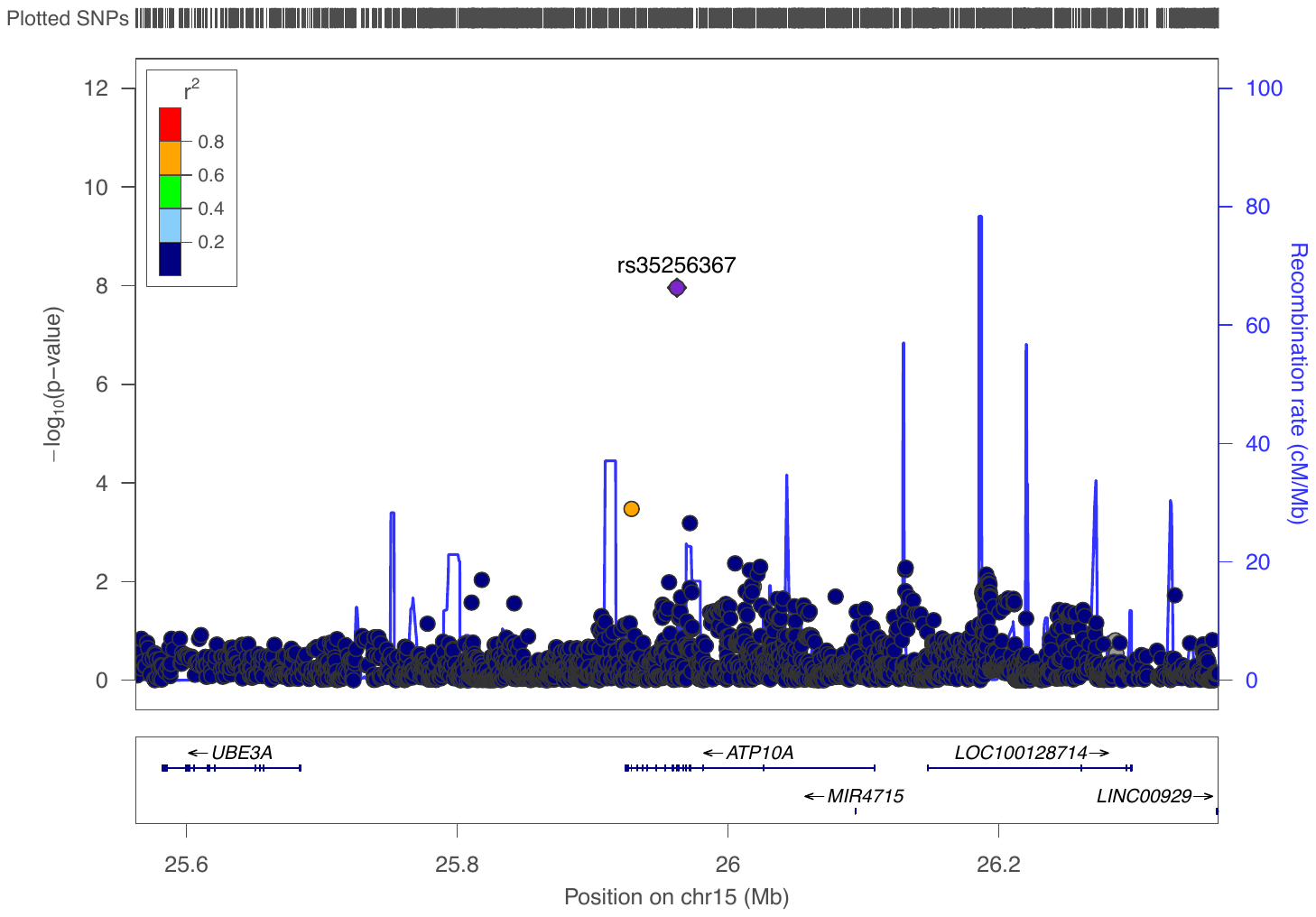

**S6. Chromosome 15**

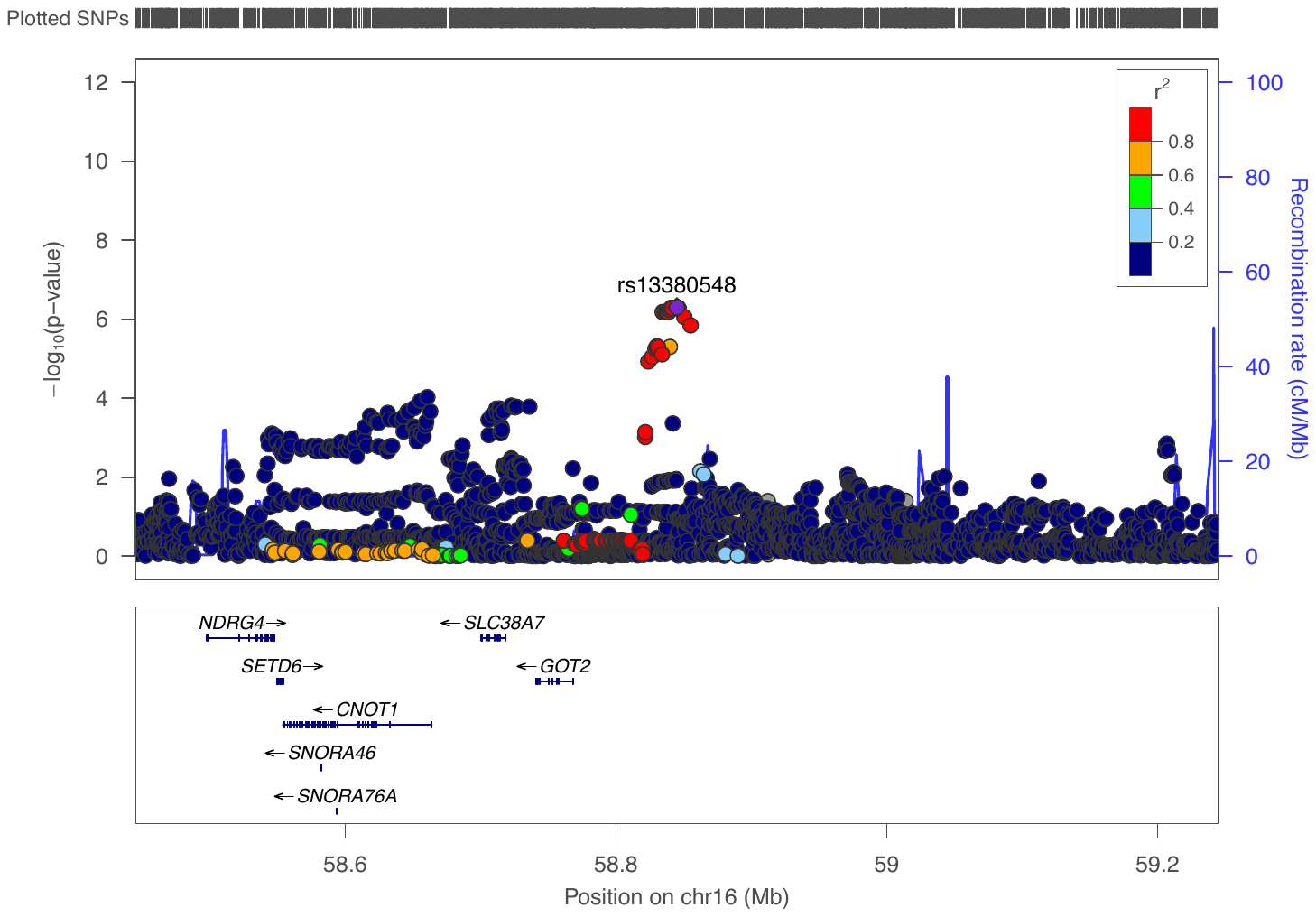

**S7. Chromosome 16**

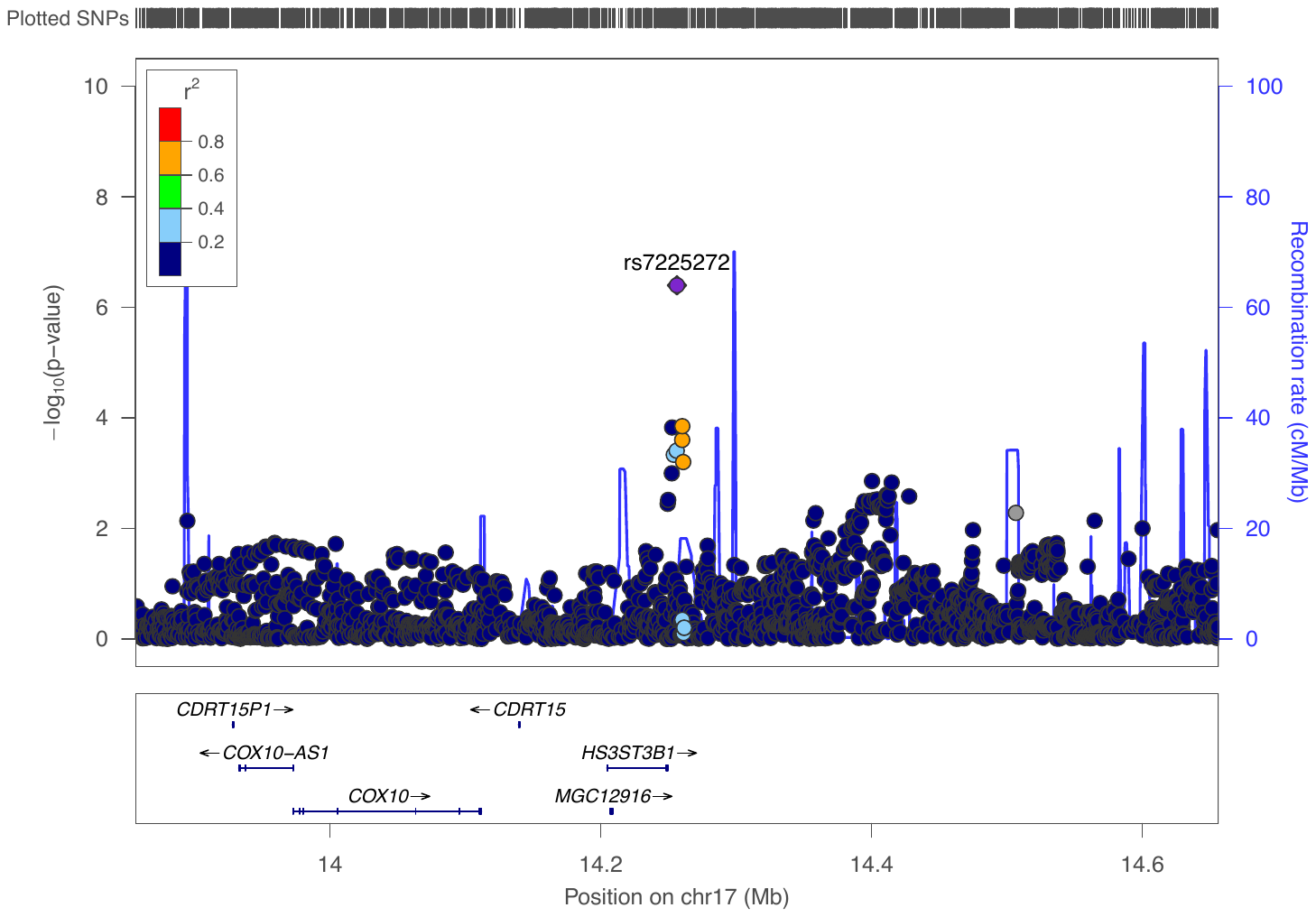

**S8. Chromosome 17**

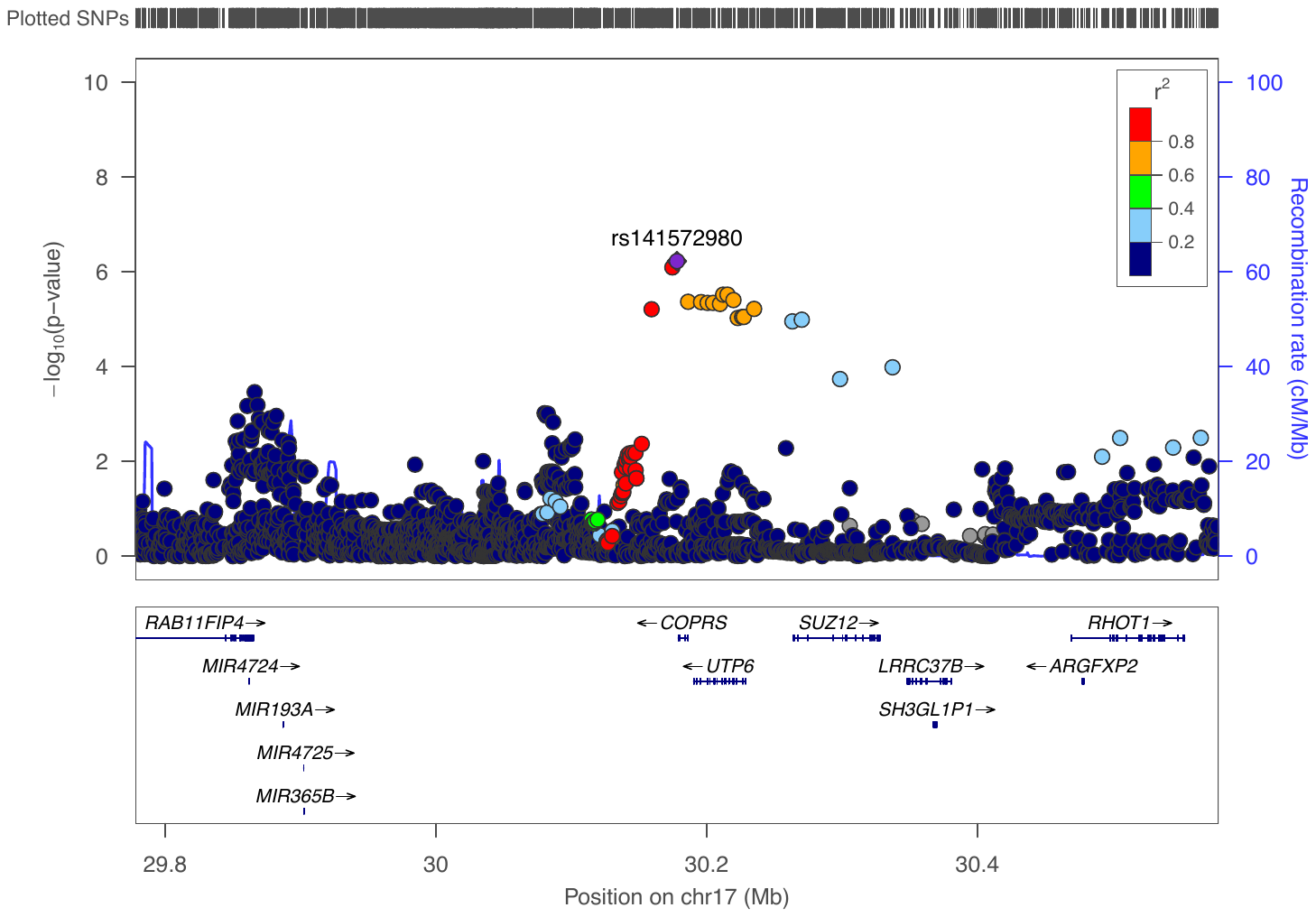

**S9. Chromosome 17**

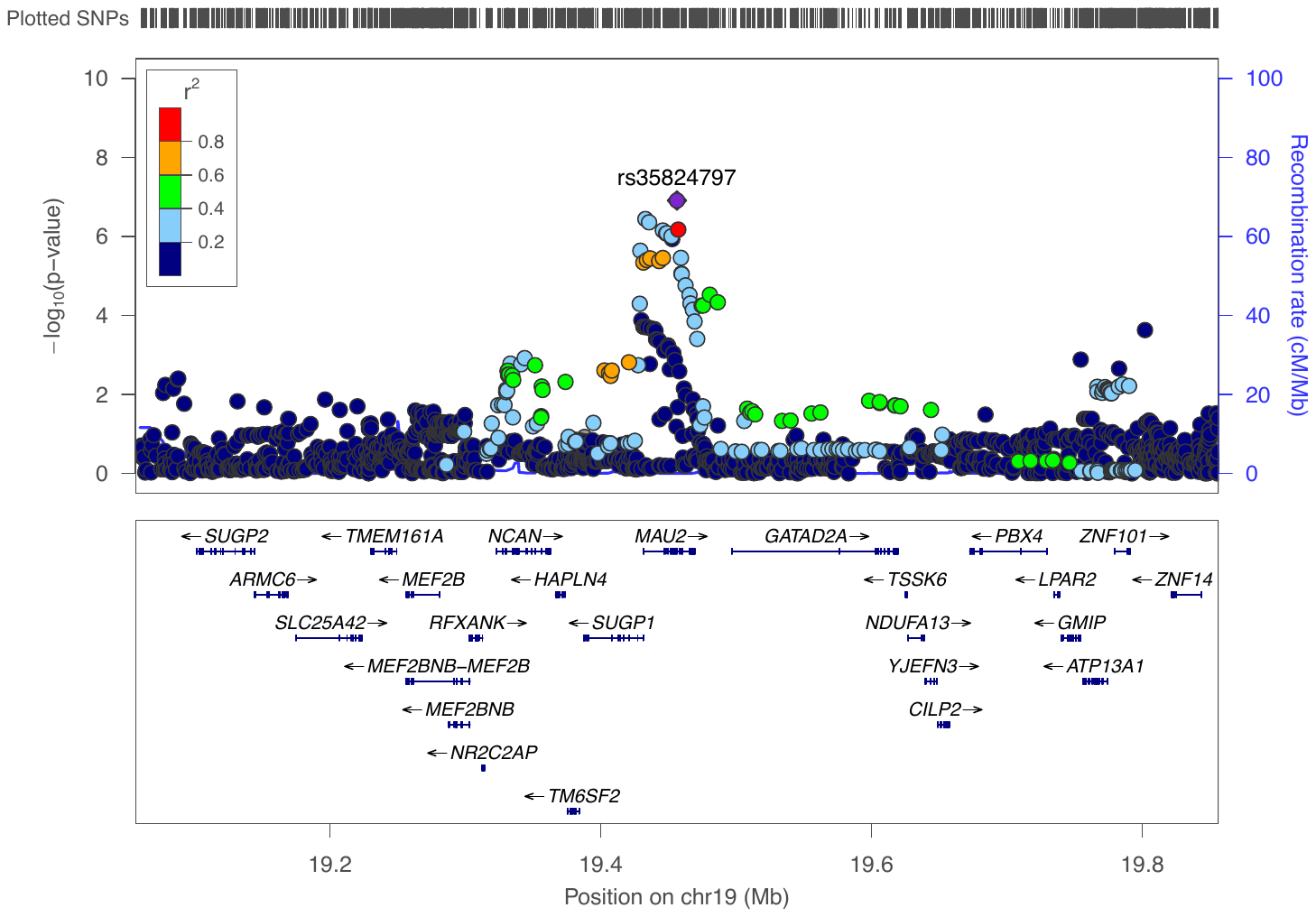

**S10. Chromosome 19**

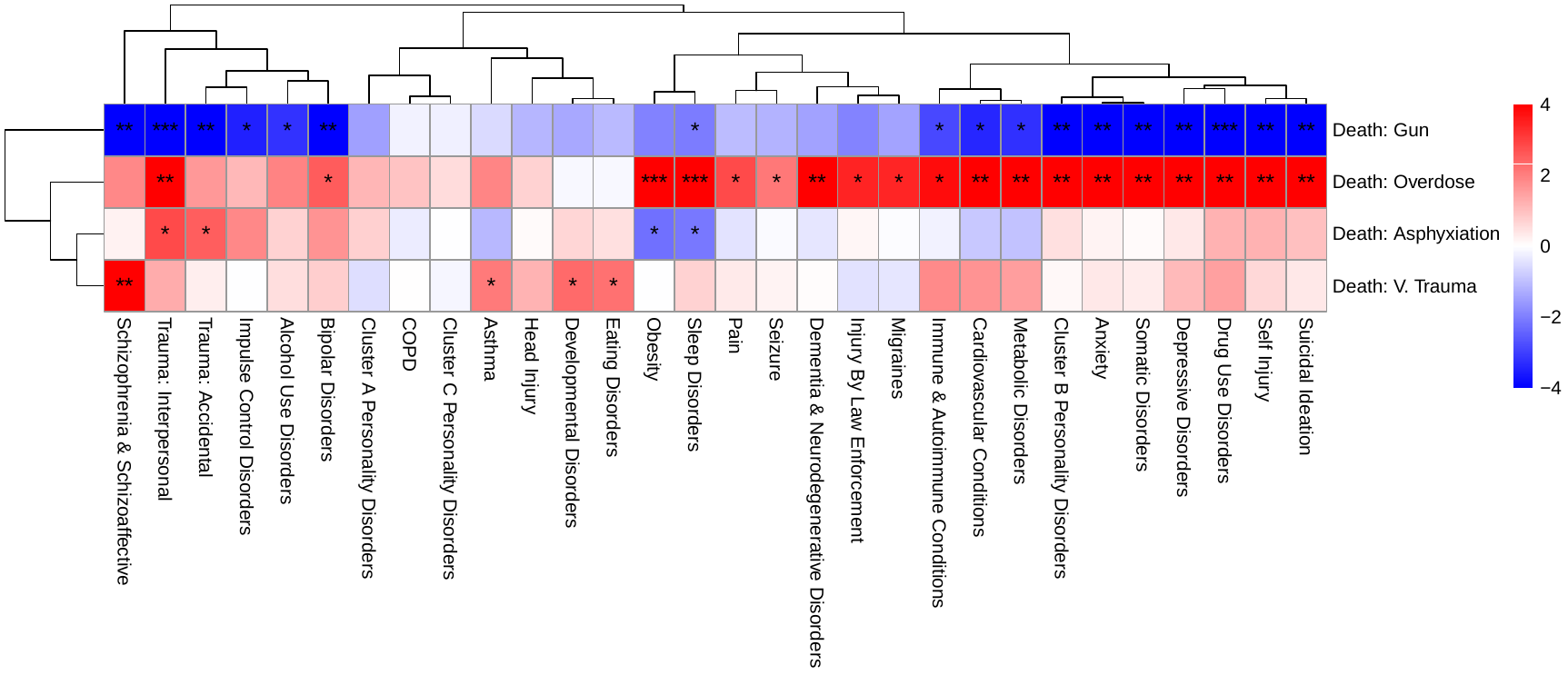

**S11.** **Diagnostic Antecedents of Specific Mode of Suicide.** Medical record diagnoses (x-axis) mapped to mode of death (y-axis) in the suicide death cases. Dendrograms based on k-means clustering of Euclidean distances are featured on both axes. Shading reflects the test statistic value as positive (red) or negative (blue). p<10^-2^ (*), p<10^-4^ (**), and p<10^-10^ (***).

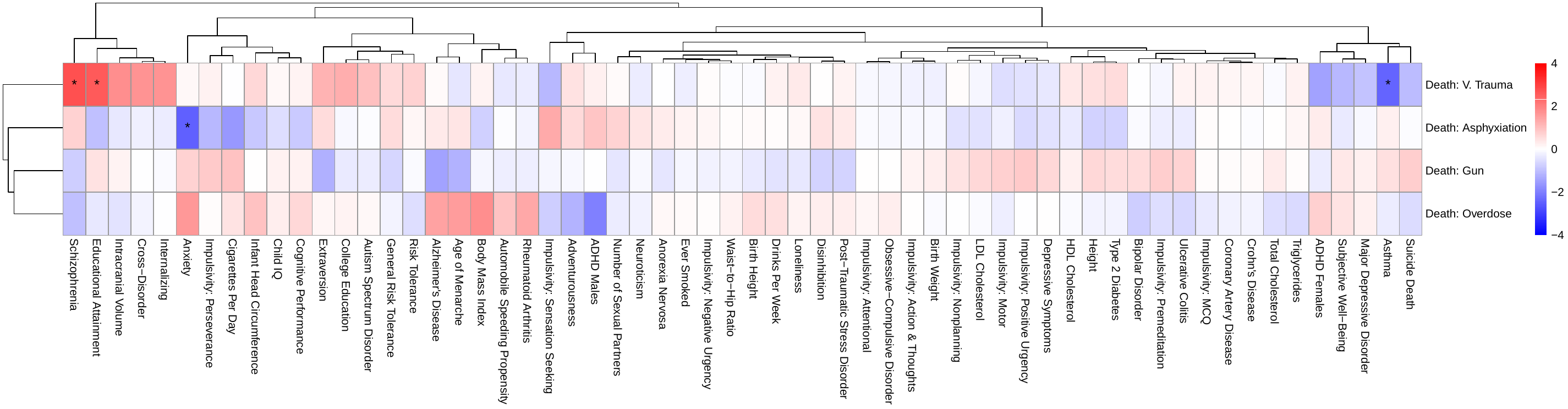

**S12. PRS X MD: All Suicide Deaths.**

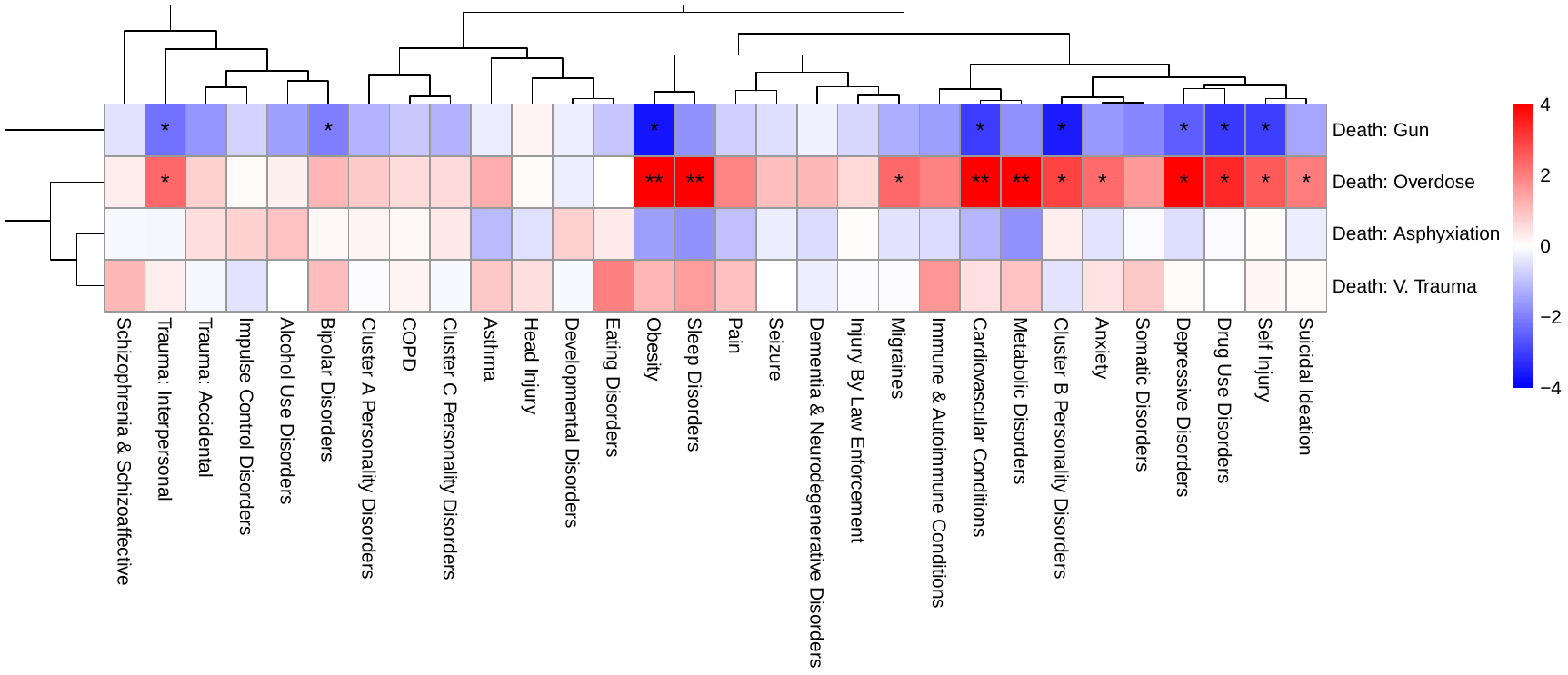

**S13. ICD X MD Plots: Female Suicide Deaths > 25 Years Of Age.**

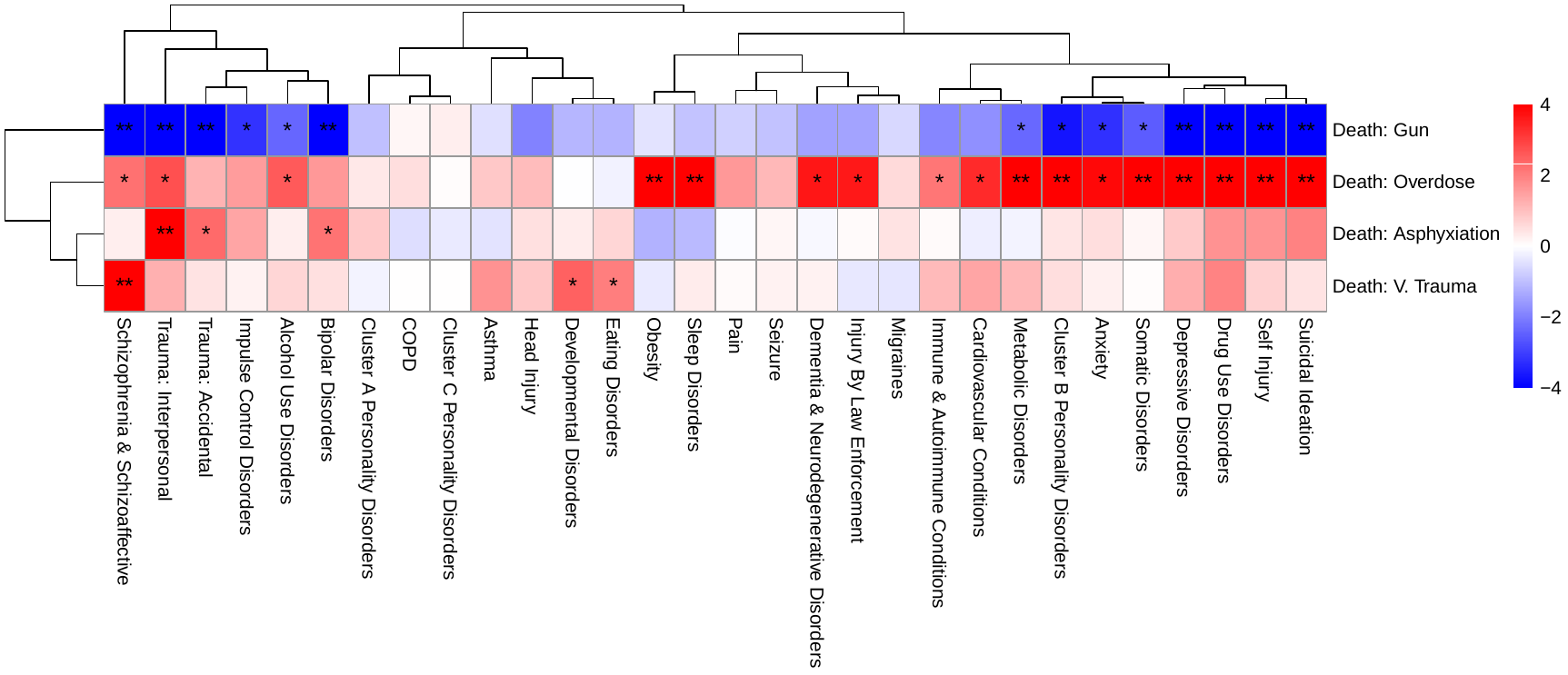

**S14. ICD X MD Plots: Male Suicide Deaths > 25 Years Of Age.**

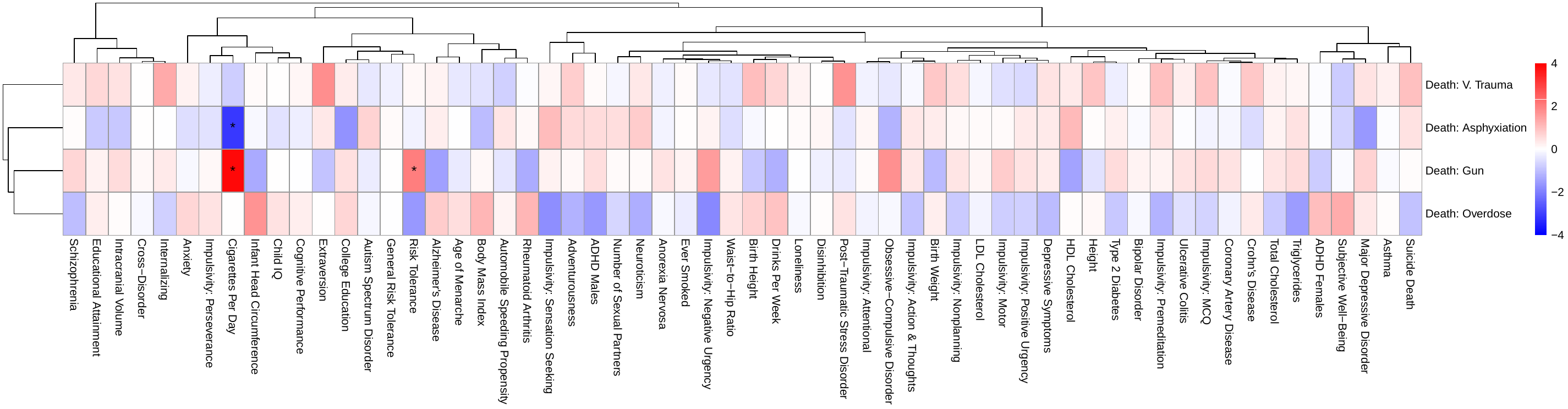

**S15. PRS X MD: Female Suicide Deaths.**

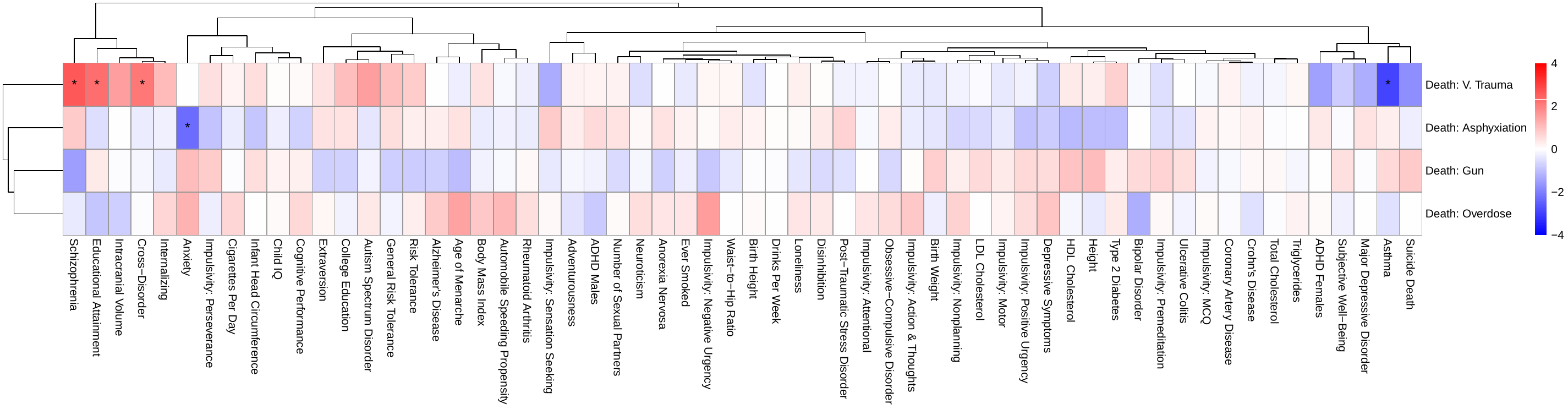

**S16. PRS X MD: Male Suicide Deaths.**
